# Guided Differentiation of Pluripotent Stem Cells into Heterogeneously Differentiating Cultures of Cardiac Cells

**DOI:** 10.1101/2023.07.21.550072

**Authors:** Erik McIntire, Kenneth A. Barr, Natalia M. Gonzales, Olivia L. Allen, Yoav Gilad

**Author notes:** Correspondence (Y.G.).

## Abstract

In principle, induced pluripotent stem cells (iPSCs) can differentiate into any cell type in the body. The challenge is to find a way to rapidly expand the dimensionality of cell types and cell states we can characterize. To address this, we developed a guided differentiation protocol to produce heterogeneous differentiating cultures of cardiac cell types (cardiac HDCs) in 16 days. Cardiac HDCs are three-dimensional, rhythmically contracting cell aggregates that harbor a temporally and functionally diverse range of cardiac-relevant cell types. We characterize cardiac HDCs from 47 iPSC lines using single-cell RNA-sequencing to identify cardiomyocytes, epicardial cells, cardiac fibroblasts, endothelial cells, and hematopoietic cells, along with both ectodermal and endodermal derivatives. This guided differentiation approach prioritizes simplicity by minimizing the reagents and steps required, thereby enabling rapid and cost-effective experimental throughput. We expect cardiac HDCs to provide a scalable cardiac model for population-level studies of gene regulatory variation and gene-by-environment interactions.

## Introduction

Induced pluripotent stem cells (iPSCs) have redefined *in vitro* cellular experimentation, offering a transformative platform for human biomedical research^1–7^. The capability to differentiate various cell types rapidly and in a controlled manner has been instrumental for population-level studies of gene regulation, which aim to clarify the relationship between genetic variation, molecular variation, and complex traits^8–11^. In particular, iPSCs make it feasible to characterize genetic variants with dynamic regulatory effects (i.e., effects that vary across different functional or temporal contexts).

Such studies have been proven quite insightful; for example, gene-by-environment interactions in human cells can explain why some individuals are at higher risk for disease when they are exposed to certain environments^7,11^, and they can reveal who is at risk for negative side effects after treatment with a drug^6^. Similarly, iPSCs can be used in time-course experiments to repeatedly measure gene regulatory activity as cells differentiate, respond to stress, or engage in other biological processes^4–6^. Still, the level of insight that can be obtained from an *in vitro* study depends on the model’s relevance to the disease, tissue, or process of interest, and in many cases, the relevant contexts are unknown^12^.

In an ideal scenario, researchers might select multiple cell types to differentiate and characterize; however, such an approach is rarely feasible for large-cohort, population-level studies. One possible solution would be to use *in vitro* tissue organoids instead of directing iPSCs to a single specific cell type. Organoids include multiple cell types and are useful for studying specific features of tissue function or for modeling diseases^13^. However, organoids are also impractical for population-level studies because, like directly differentiated iPSCs, they are difficult and time-consuming to establish. Because the optimal culture conditions for different cell types within an organoid are often conflicting, it can be challenging to maintain multiple cell types in the same medium^14,15^. As a result, organoid culture tends to be inefficient and takes weeks to months to establish. Further, organoids can be difficult to dissociate to the single-cell level for sequencing and characterization of regulatory phenotypes^16,17^.

These challenges propel our exploration of ‘guided’ differentiation, an approach that aims to balance protocol efficiency and cellular diversity. We call our approach ‘guided’ differentiation, because unlike ‘directed’ differentiation, which pushes cells toward a single cell type, we gently bias iPSCs towards a broader lineage, aiming to maintain the greatest degree of cellular diversity possible. Focusing on cardiac cell types, we adapted methods used in embryoid body and cardiac organoid generation to guide iPSCs toward the cardiac mesoderm lineage, resulting in heterogeneous differentiating cultures of cardiac cell types (cardiac HDCs; **Figure 1A**). Cardiac HDCs are tailored for population-level studies of dynamic gene regulation. In addition to harboring a diversity of cardiac-relevant cell types, they are designed for efficiency, requiring minimal protocol steps and reagents, and allowing for high experimental throughput. Here, we provide a detailed protocol to establish cardiac HDCs using guided differentiation. We anticipate that cardiac HDCs will have many applications in population-level studies; examples include associating genetic variants with cardiovascular traits, studying cell-type-specific responses to drugs and treatments, and exploring gene-by-environment interactions.

**Figure 1.**
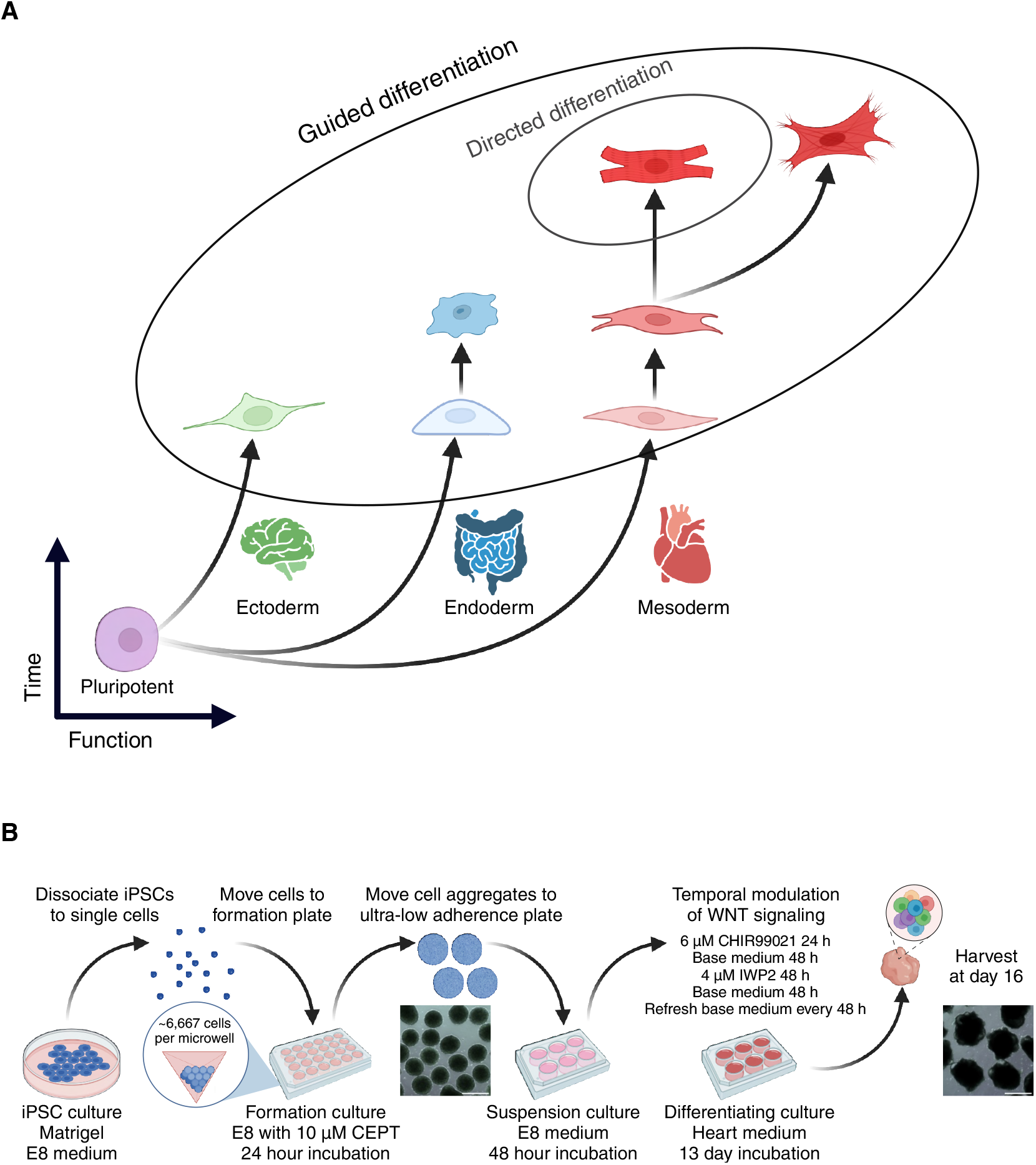
Guided differentiation of pluripotent stem cells to generate cardiac HDCs. (A) Guided differentiation is designed to harbor cells at multiple differentiation stages across all three germ layers while maintaining predominantly heart-associated cell types in the case of cardiac HDCs. (B) 16-day guided differentiation protocol. IPSCs are dissociated to form three-dimensional aggregates using an AggreWell™800 24-well plate. After formation, iPSC aggregates are moved to a suspension culture for 48 h in E8 medium. Cell aggregates are differentiated using temporal modulation of Wnt signaling with CHIR99021 and IWP2. Scale bars 500 μm. Abbreviations: E8: Essential 8™ medium. CEPT: Chroman 1, emricasan, polyamines, and trans-ISRIB. Base heart medium: RPMI 1640 Medium, GlutaMAX™ Supplement, HEPES, 2% v/v B-27 Supplement, minus insulin.

## Results

We performed guided differentiation (**Figure 1B**) using 47 iPSC lines to generate cardiac HDCs. First, we formed iPSCs into three-dimensional aggregates measuring ∼300 μm in diameter, the optimal size for inducing pluripotent stem cells towards the cardiac lineage^18,19^. We cultured iPSC aggregates in the formation plate using Essential 8™ medium (E8) with 10 μM CEPT for 24 hours and transferred the aggregates to ultra-low attachment plates for a full 48 hours with only E8^20^. We then biased iPSC aggregates towards the cardiac lineage using temporal Wnt modulation^21,22^. We exchanged E8 for ‘heart medium’ (RPMI-1640 with a 2% v/v concentration of B-27 supplement, *sans* insulin) plus the Wnt activator CHIR99021 (‘Chiron’) at a final concentration of 6 μM. After 24 hours, we exchanged Chiron+ heart medium for base heart medium. We refreshed the heart medium 48 hours later, this time adding IWP2, a Wnt inhibitor, at a final concentration of 4 μM. After 48 hours, we replaced IWP2+ heart medium with base heart medium and proceeded to refresh with base heart medium every 48 hours until harvest on day 16. Unlike standard cardiac differentiation, no insulin is added during guided differentiation. We collected cells at day 16 using the 10x Genomics platform for sequencing on an Illumina NovaSeq X.

After filtering and normalizing the data, we performed principal component analysis with 5,000 highly variable features and used the top 50 principal components for graph-based unsupervised clustering. We classified cardiac HDC cell types using marker gene expression and differential expression analysis, annotating a total of 14 different cell types representing all three germ layers with both progenitor and terminally differentiated cell types (**Figure 2**). Within cardiac HDCs, individual cells differentiate at dissimilar rates, permitting diversity with respect to function (cell type) and time (cell differentiation stage). The inclusion of early progenitor cells, in addition to differentiated cells, provides developmental trajectories for detecting transient genetic effects that typically can only be captured across several collections during time-course experiments^4,5^. Guided differentiation consistently generated this wide diversity of cell types, including cardiomyocytes, across all 47 iPSC lines (**Figure Supplement 1**).

**Figure 2.**
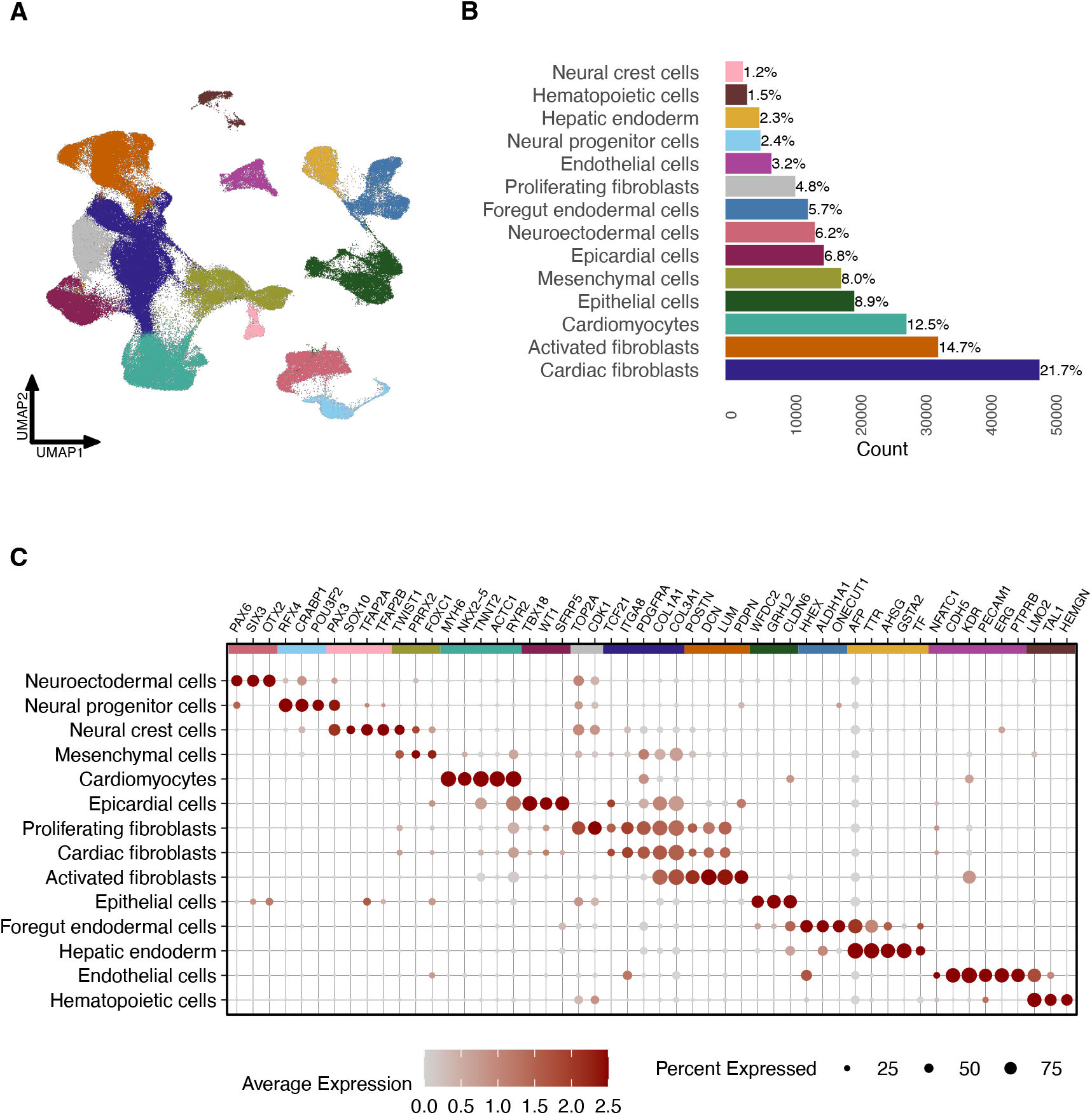
Single-cell RNA-sequencing characterization of cardiac HDCs. (A) Uniform manifold approximation and projection (UMAP) plot of 222,438 cells from cardiac HDCs generated from 47 iPSC lines. (B) Percentages of 14 distinct cell types within cardiac HDCs, as identified by manual annotation based on marker gene expression. (C) Dot plot depicting the relative expression of marker genes across cell types, where dot size reflects the proportion of cells expressing each gene within a cluster and color intensity represents gene expression levels.

We classified cardiomyocytes per expression of sarcomere genes (*TNNT2, MYH6, ACTC1*) along with the ion channel gene *RYR2* and the transcription factor *NKX2-5*^23,24^. We annotated the epicardial cell cluster using *SFRP5, WT1*, and *TBX18*^25^. We identified three clusters of fibroblasts based on expression of *TCF21, ITGA8, PDGFRA, COL1A1*, and *COL3A1*^26–30^. Cardiac fibroblasts were marked by this core set of genes; proliferating fibroblasts additionally expressed *CDK1* and *TOP2A*^31,32^. Activated fibroblasts were characterized by the core gene set, along with elevated expression of extracellular matrix components (*POSTN, DCN*, and *LUM)*, coupled with reduced expression of *TCF21*^33^ and cluster-specific *PDPN* expression, a marker for fibroblast activation^34^. Finally, we designated a cell cluster of mesenchymal cells based on expression of *TWIST1* and *PRRX2*, and *FOXC1*^35,36^.

In addition to these mesoderm-derived cell types, cardiac HDCs also harbor cell types from endodermal and ectodermal lineages. Endodermal cells support cardiac cell differentiation^37,38^. We classified two endoderm populations: foregut endoderm based on expression of *HHEX, ONECUT1*, and *ALDH1A1*^39–41^, and hepatic endoderm per expression of *AFP, TTR, AHSG*, and *TF*^42,43^. We characterized epithelial cells using expression of *GRHL2, WFDC2*, and *CLDN6*^44–46^. Lastly, we characterized three cell clusters from the ectodermal lineage: neuroectodermal (*OTX2, SIX3, PAX6*^47^) and neural progenitor cells (*POU3F2, CRABP1, RFX4*^48–50^) as well as neural crest cells (*SOX10, PAX3, TFAP2A*, and *TFAP2B*^51–54^).

Cardiac HDCs also exhibit cell types related to vasculogenesis and hematopoiesis, i.e., endothelial and hematopoietic cells. We classified endothelial cells per expression of *CDH5, KDR, PECAM1, ERG*, and *PTPRB*^55^. A small proportion of these cells show strong expression of *NFATC1*, a canonical marker for endocardial cells^56^. Expression of *LMO2, TAL1*, and *HEMGN* revealed a cluster of hematopoietic cells that we subset into smaller groups (**Figure 3**)^57–59^. Sub-clustering revealed that the hematopoietic population primarily consists of erythrocytes (*HBA2, SLC4A1, ALAS2, GYPA*^60^) and megakaryocytes (*PF4, PPBP, TMEM40, GP1BA*^61^), with small populations of monocytes (*PTPRC, LYZ, CSF1R, SPI1*^62^) and hematopoietic progenitor-like cells (*ZNF608, TGFBR3, SOX6, YAP1*^63–66^). This feature of cardiac HDCs parallels a recent observation in blood-generating, heart-forming organoids that support concurrent development of cardiomyocyte, endothelial, and hematopoietic cell types (among others)^67^.

**Figure 3.**
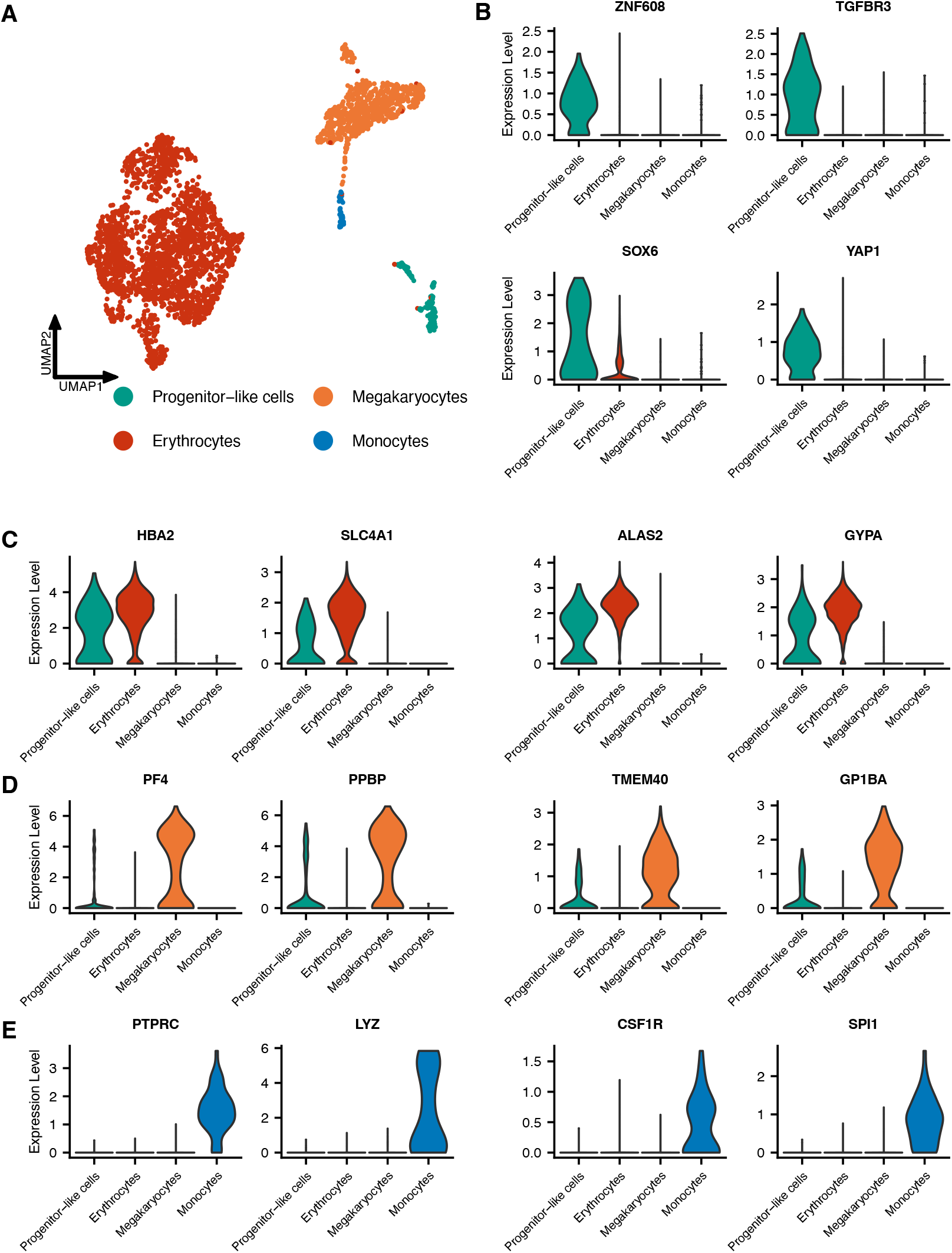
Sub-clustering of hematopoietic cells reveals distinct populations. (A) UMAP plot of 3,340 sub-clustered hematopoietic cells. (B–E) Violin plots showing expression of marker genes for progenitor-like cells (B), erythrocytes (C), megakaryocytes (D), and monocytes (E).

We next investigated similarity between the transcriptional profiles of cells from cardiac HDCs and their *in vivo* counterparts using popular vote (popV) automated classification^68^, trained using reference datasets from Tabula Sapiens^69,70^. PopV classifies cell types based on the consensus of eight prediction models. We applied popV label transfer to classify cardiomyocytes, hepatic endoderm, endothelial cells, and hematopoietic cells in cardiac HDCs using adult heart, liver, endothelium, and immune datasets, respectively (**Figure 4**). To ensure label transfer accuracy, we retained only query cells where at least six of eight prediction models agreed (a ‘high consensus’ score). The results corroborate our manual cell annotation and demonstrate that cells in cardiac HDCs transcriptionally resemble those in corresponding adult tissues.

**Figure 4.**
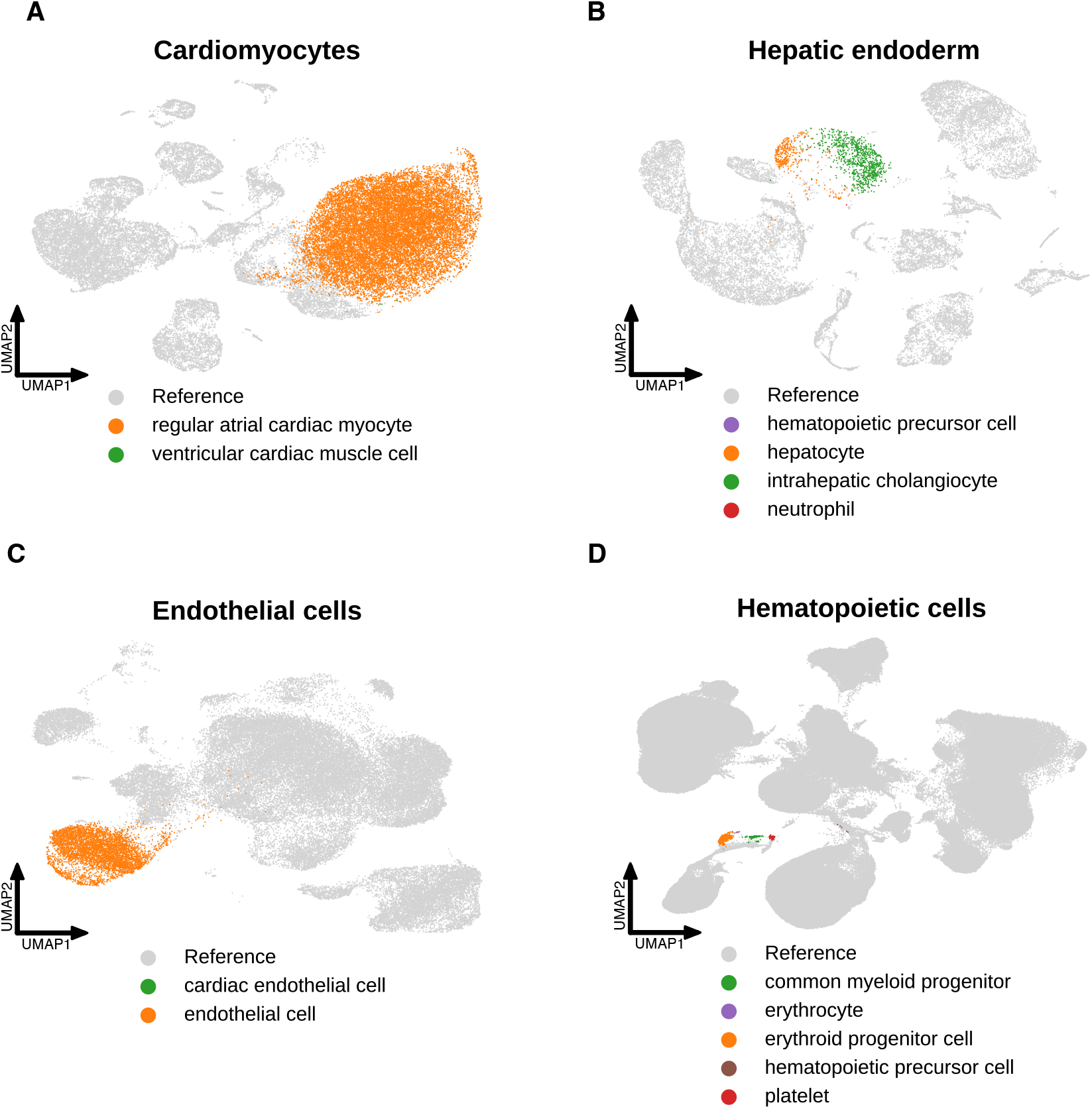
High consensus popV label transfer demonstrates transcriptional similarity between cardiac HDC cell types and their adult tissue counterparts. UMAP plots showing the mapping of cardiac HDC cell populations (colored) onto adult reference datasets (gray): cardiomyocytes to heart (A), hepatic endoderm to liver (B), endothelial cells to endothelium (C), and hematopoietic cells to immune cells (D).

## Discussion

We present cardiac HDCs, a human cell culture model that combines cardiac cell diversity with labor and reagent efficiency to address an unmet need in functional genomics. Population-level genomic studies typically involve large, complex datasets, and they frequently require access to multiple different cell types. Standard embryoid body and cardiac organoid approaches offer cell type diversity, but in most cases, complexity and inefficiencies render them impractical for population-level studies of dynamic gene regulation. By contrast, guided differentiation produces a diverse spectrum of cardiac-relevant cell types within the short period of 16 days, while still recapitulating the cellular complexity observed in mature cardiac organoids. Guided differentiation also permits cells to differentiate at varying (asynchronous) rates, such that a single collection of cardiac HDCs comprises cells at various states along a developmental continuum. Thus, cardiac HDCs make it possible to capture transient genetic effects at greater temporal resolution than any time-course study could reasonably achieve.

The ability to grow multiple cardiovascular cell types and cell states in the same dish obviates the need for complex differentiation protocols and allows for greater control over confounding variables that might mask genetic effects on gene expression. We expect that the reduced time and labor needed for guided differentiation (relative to standard organoids and differentiation time-course studies) will enable dynamic population-level studies at scale. For example, guided differentiation may facilitate the identification of gene-by-environment interactions and molecular quantitative trait loci (e.g., expression QTLs). Moreover, by capturing transient genetic effects exclusive to early development – namely, effects that may not be detectable in terminal cell types or adult tissues^4,5^ – guided differentiation could be leveraged to interpret as-yet unexplained genetic associations with disease^71^.

## Acknowledgements

We thank all members of the Gilad lab for their support. This work was completed in part with resources provided by the University of Chicago’s Research Computing Center. We thank the University of Chicago Functional Genomics Core Facility for their assistance with sequencing the libraries. Figures 1A and 1B were created using BioRender.com.

## Author contributions

Y.G. conceived the study and supervised the project. E.M. developed the differentiation method and performed the differentiation experiments. E.M., O.L.A., and K.A.B performed single cell RNA-seq library preparation. E.M. analyzed and interpreted the data with support from K.A.B. E.M. drafted the manuscript with review and editing provided by Y.G and N.M.G.

## STAR METHODS

### Key resources table

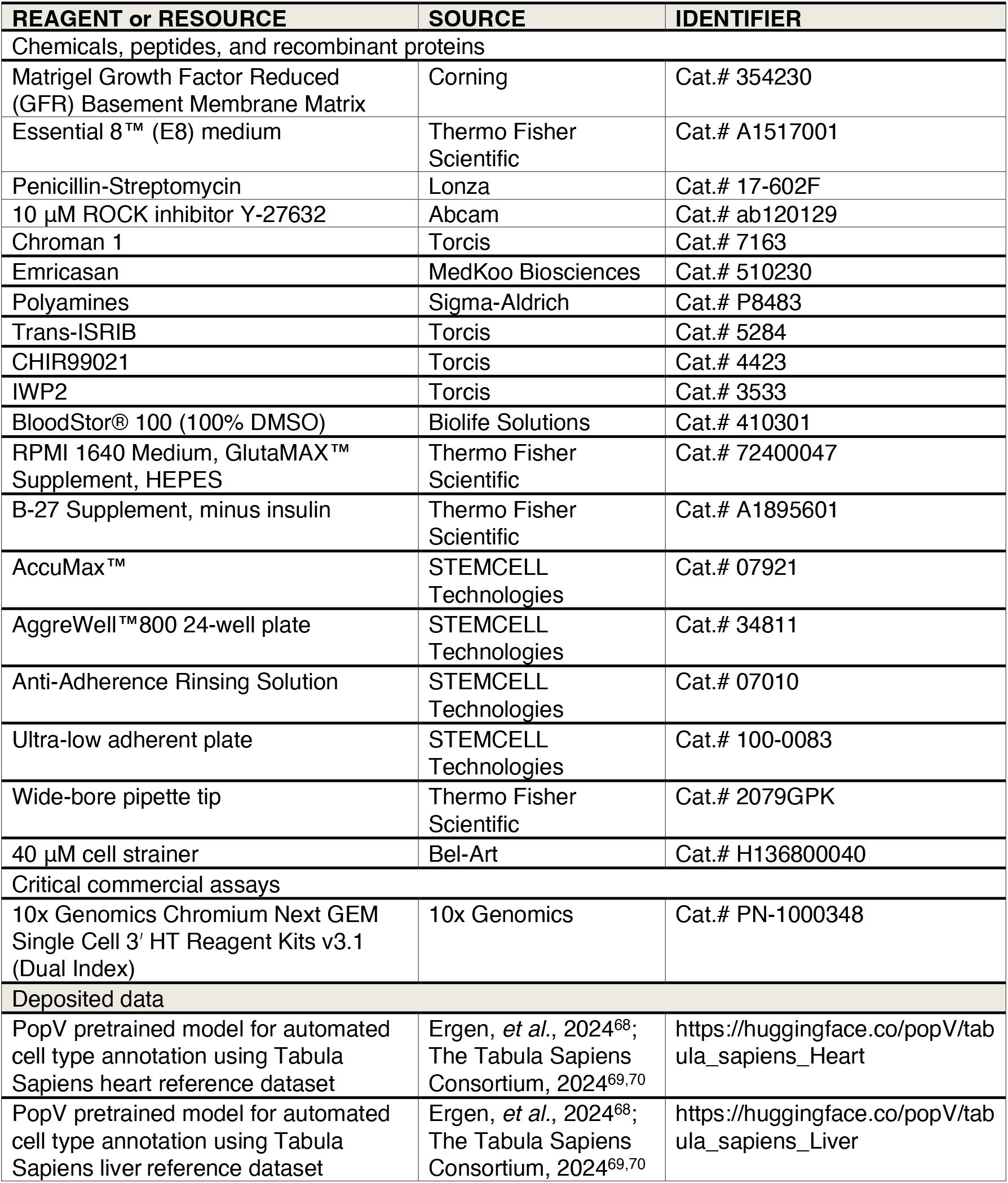

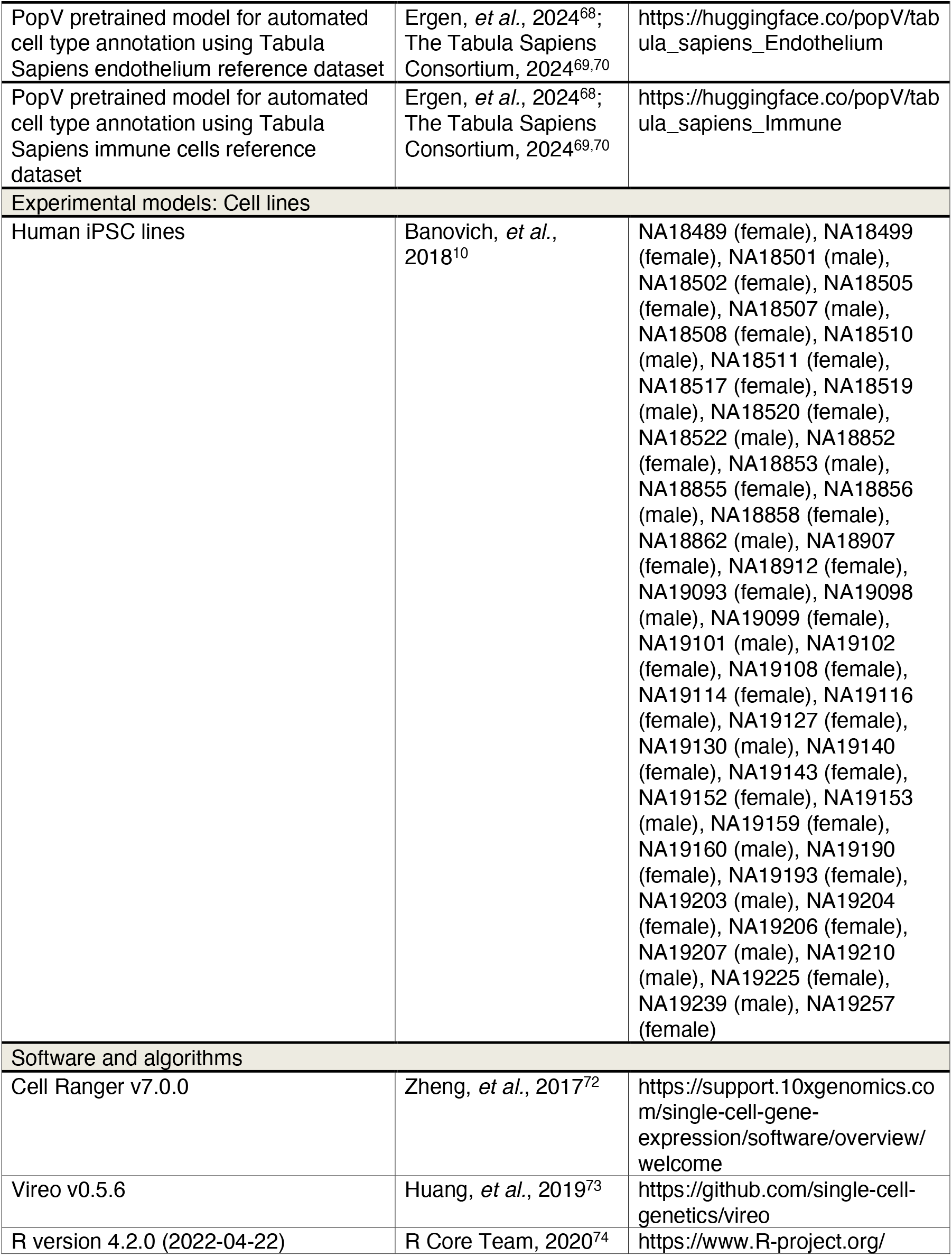

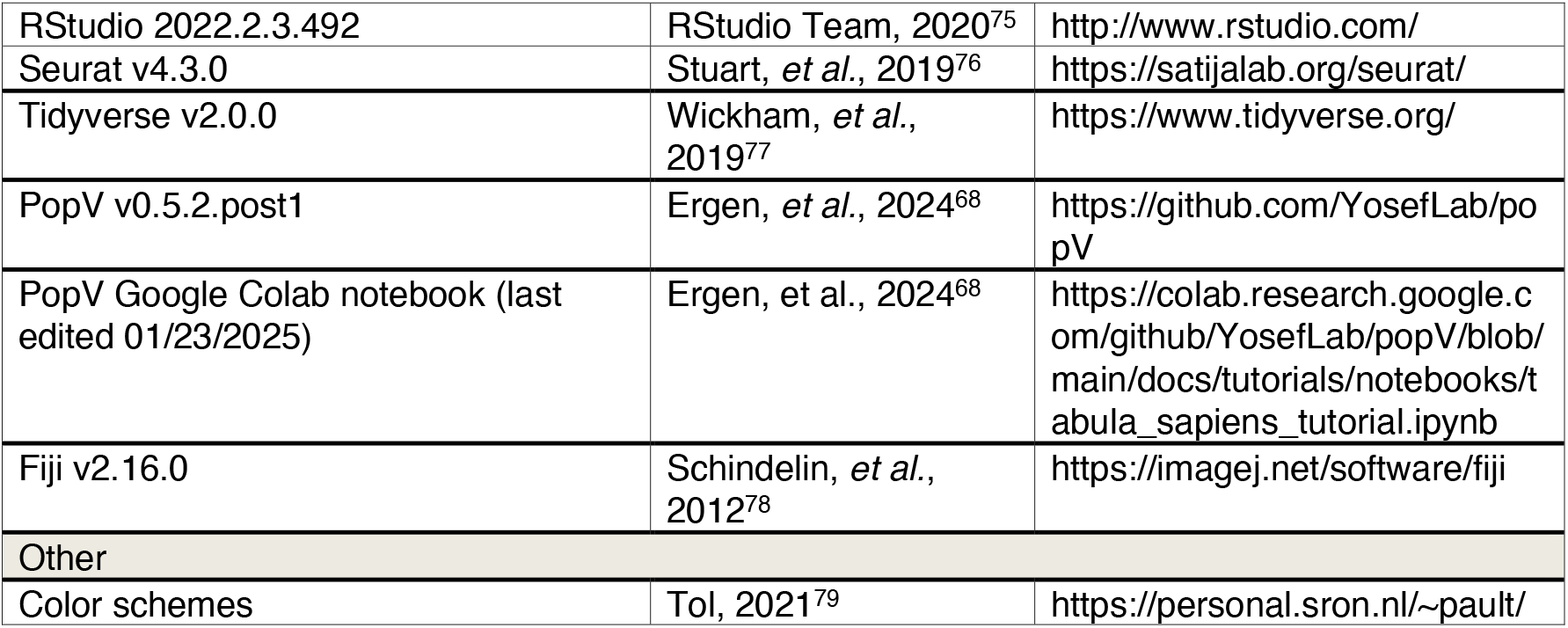

## Resource availability

### Lead contact

Further information and requests for resources should be directed to and will be fulfilled by the lead contact, Dr. Yoav Gilad.

## Materials availability

This study did not generate any new reagents.

## Data and code availability

Code for data analysis available at: https://github.com/erikmcintire/Cardiac_HDCs Sequencing data will be made public concurrent with publication.

## Method details

### Samples

We used 49 iPSC lines (47 after quality control and filtering, 17 male and 30 female) from unrelated Yoruba individuals from Ibadan, Nigeria (YRI). These iPSC lines were reprogrammed from lymphoblastoid cell lines (LCLs) and were characterized and validated previously^10^. We confirmed cell line identities using genotype data generated by the HapMap project from the original LCL lines^80^.

### IPSC maintenance

We maintained iPSC lines using Matrigel Growth Factor Reduced (GFR) Basement Membrane Matrix (354230, Corning) with Essential 8™ (E8) medium (A1517001, Thermo Fisher Scientific) and Penicillin-Streptomycin (17-602F, Lonza) in an incubator at 37°C and 5% CO_2_. At roughly 80% confluency (approximately every 3-5 days), we passaged cell cultures using a dissociation reagent (0.5 mM EDTA, 300 mM NaCl in PBS) and seeded iPSCs with 10 μM ROCK inhibitor Y-27632 (ab120129, Abcam).

### Guided differentiation

We formed and harvested cardiac HDCs across six batches, each consisting of six to ten iPSC lines, balanced by sex and passage number. We performed guided differentiation on iPSCs by forming three-dimensional aggregates and applying temporal Wnt modulation^21,22^. Small molecules CHIR99021 (4423, Torcis) and IWP2 (3533, Torcis) were dissolved in BloodStor® 100 (100% DMSO) (410301, Biolife Solutions) at stock concentrations of 10 mM and 4 mM, respectively. Stock solutions were stored at -20 °C and remade at least every 3 months. We formed iPSC aggregates using an AggreWell™800 24-well plate (34811, STEMCELL Technologies), with each well coated with Anti-Adherence Rinsing Solution (07010, STEMCELL Technologies). On day 0, we dissociated iPSCs (using 0.5 mM EDTA, 300 mM NaCl in PBS) and seeded them onto the plate with E8 medium and 10 μM CEPT, i.e., chroman 1 (7163, Torcis), emricasan (510230, MedKoo Biosciences), polyamines (P8483, Sigma-Aldrich), and trans-ISRIB (5284, Torcis)^20^. Aggregates were formed at approximately 300 μm diameter by seeding at a density of 2 million cells per well, as described previously by Branco *et al*. (2019)^19^. We centrifuged the plate at 100 *g* for 3 min to aggregate cells, and cells were maintained in the plate for 24 h. After 24 h, we transferred the aggregates to a suspension culture using ultra-low adherent plates (100-0083, STEMCELL Technologies) with 4 mL of medium per well. First, cells were placed in E8 medium for 48 h; note that the cells were maintained in the initial E8 medium throughout the entire 48-hour duration with no replenishment. On day 3, we switched the cell culture medium to ‘heart medium’ plus 6 μM CHIR99021. Heart medium consists of RPMI 1640 Medium, GlutaMAX™ Supplement, HEPES (72400047, Thermo Fisher Scientific), 2% v/v B-27 Supplement, minus insulin (A1895601, Thermo Fisher Scientific), and Penicillin-Streptomycin. After 24 h, on day 4, we changed the medium to base heart medium for 48 h. On day 6, we changed the medium to heart medium plus 4 μM IWP2 for 48 h. From day 8 onward, cells were maintained in base heart medium, and we refreshed medium every 48 h (days 8, 10, 12, 14). On day 15, we performed a final base heart medium refresh (containing 0.05% DMSO, which served as a vehicle control in a separate experimental study). We harvested cardiac HDCs on day 16. We imaged cardiac HDCs using an EVOS^®^ XL Core Imaging System and processed images with Fiji (v2.16.0)^78^.

### Aggregate dissociation

We collected day 16 aggregates and dissociated them by treating with room temperature AccuMax™ (07921, STEMCELL Technologies) followed by incubation at 37 °C for 10 min. Following incubation, we pipetted aggregates up-and-down for 30 sec with a p1000 wide-bore pipette tip (2079GPK, Thermo Fisher Scientific). Aggregates were then incubated for an additional 5 min at 37 °C. We repeated the pipetting every 5 min until aggregates were dissociated, at which point we added 5 mL of 4 °C base heart medium and centrifuged cells for 100 *g* for 3 min. We resuspended cells in 1 mL 4 °C base heart medium and strained cells through a 40 μm cell strainer (H136800040, Bel-Art). We combined cells from each iPSC line in even proportions and centrifuged cells at 100 *g* for 3 min. We then resuspended cells in 4 °C base heart medium at a concentration of approximately 1.6 million cells per mL.

### Single-cell RNA sequencing analysis

We collected cells from cardiac HDCs for scRNA-seq using the 10x Genomics Chromium Next GEM Single Cell 3’ HT Reagent Kits v3.1 (Dual Index) (PN-1000348, 10x Genomics). For each batch, we used two lanes of a 10x chip, recovering around 20,000 cells per lane (∼40,000 cells per batch). We sequenced the libraries using an Illumina NovaSeq X at the University of Chicago Functional Genomics Core Facility, yielding 35,897 mean reads per cell. We aligned samples to the human genome (GRCh38) using Cell Ranger (v7.0.0)^72^ and assigned cells to individuals using Vireo (v0.5.6)^73^. Two iPSC lines were removed from batch 1 due to cross-contamination. Cell line NA19114 was mistakenly collected twice (in batches 1 and 5) due to mislabeling. We analyzed count data in R (v4.2.0)^74^ / RStudio (v2022.2.3.492)^75^ using Seurat (v4.3.0)^76^ with Tidyverse (v2.0.0)^77^. We merged all Seurat objects and performed cell filtering by removing genes expressed in fewer than 3 cells, removing cells classified as doublets or unassigned by Vireo, and removing cells with fewer than 1,500 unique genes. After filtering, 33,948 genes were retained across 222,438 cells from 47 unique iPSC lines for downstream analysis. We performed log-based normalization of gene expression. We then centered and scaled counts for the 5,000 most highly variable genes. During scaling, we regressed out cell cycle phase differences to minimize variability among proliferating cells while preserving phase differences between cycling and non-cycling cells, thereby maintaining distinctions between progenitor and terminally differentiated populations. We applied principal components analysis (PCA) and used the top 50 principal components for uniform manifold approximation and projection (UMAP) embedding. We computed k-nearest neighbors and then performed unsupervised clustering at a resolution of 0.12, yielding 14 cell clusters for differential expression analysis using the Wilcoxon Rank Sum test. We annotated cell clusters based on expression of key marker genes. For figures 2 and 3, we shaded cell clusters using color schemes adapted from Paul Tol^79^.

### Hematopoietic cells sub-cluster analysis

We subset the hematopoietic cells as their own Seurat object and re-normalized using the same approach described above. We performed unsupervised clustering at a resolution of 0.075 to yield 4 clusters for differential expression analysis. We annotated cell clusters based on marker gene expression.

### Automated cell classification

We used popV (v0.5.2.post1)^68^ to perform automated cell annotation of cardiac HDC cell types using reference datasets from Tabula Sapiens^69,70^. We subset cardiac HDC cell populations of cardiomyocytes, hepatic endoderm, endothelial cells, and hematopoietic cells and performed popV label transfer using pretrained models from adult heart, liver, endothelium, and immune reference datasets, respectively. We performed analysis using the popV Google Colab Notebook (last edited 01/23/2025), provided in their GitHub repository. Of the eight default prediction models used for label transfer, we retained only query cells for which at least six of the eight prediction models agreed (a “high consensus” score).

**Figure supplement 1.**
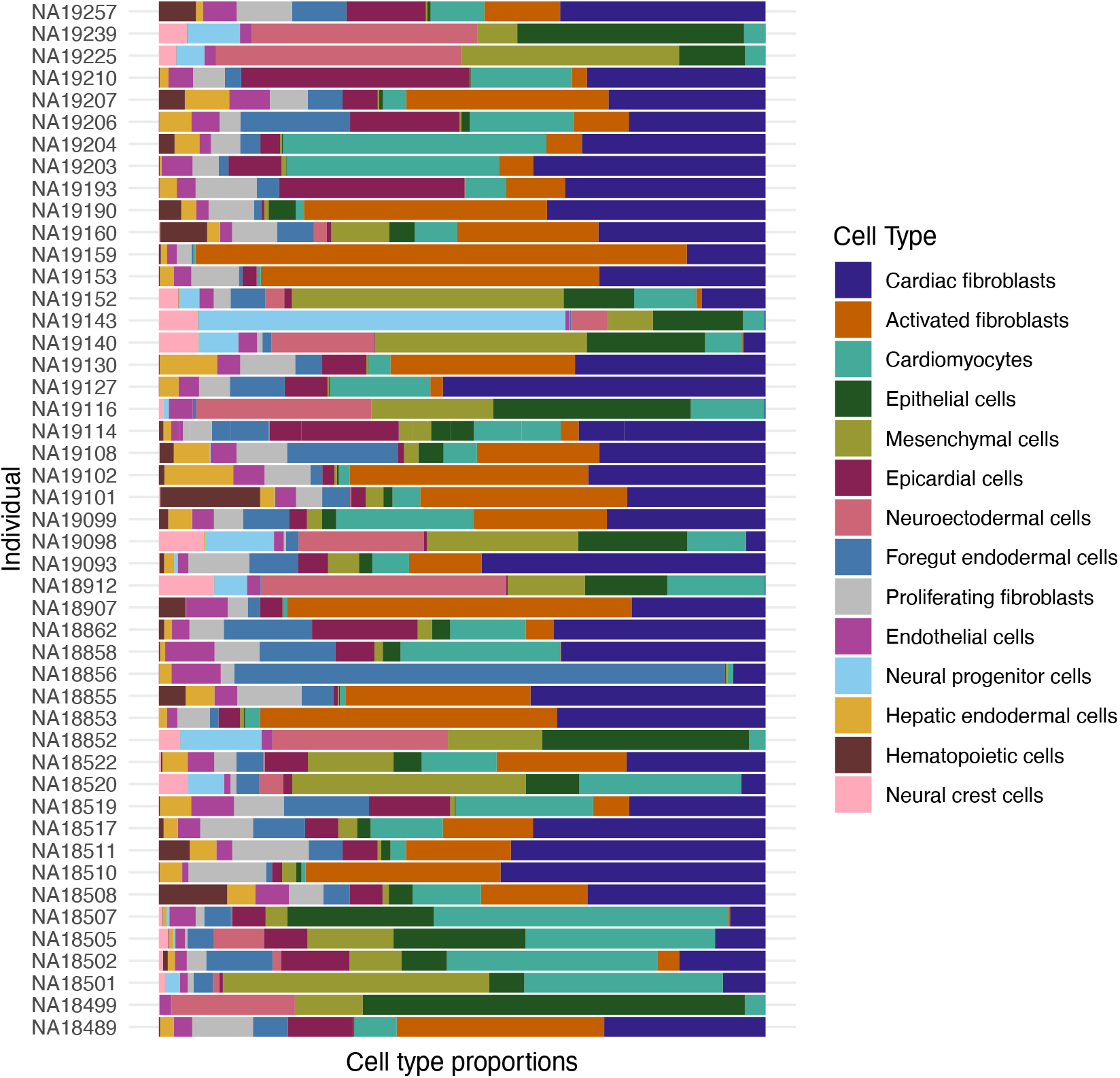
Stacked bar plot of cell type proportions across 47 iPSC lines. Guided differentiation consistently produced cardiac HDCs containing diverse cell types, including cardiomyocytes in every iPSC line.

## References

1. Takahashi, K. et al. Induction of Pluripotent Stem Cells from Adult Human Fibroblasts by Defined Factors. Cell 131, 861–872 (2007).

2. Shi, Y., Inoue, H., Wu, J. C. & Yamanaka, S. Induced pluripotent stem cell technology: a decade of progress. Nat. Rev. Drug Discov. 16, 115–130 (2017).

3. Rowe, R. G. & Daley, G. Q. Induced pluripotent stem cells in disease modelling and drug discovery. Nat. Rev. Genet. 20, 377–388 (2019).

4. Strober, B. J. et al. Dynamic genetic regulation of gene expression during cellular differentiation. 5 (2019).

5. Elorbany, R. et al. Single-cell sequencing reveals lineage-specific dynamic genetic regulation of gene expression during human cardiomyocyte differentiation. PLOS Genet. 18, e1009666 (2022).

6. Knowles, D. A. et al. Determining the genetic basis of anthracycline-cardiotoxicity by molecular response QTL mapping in induced cardiomyocytes. eLife 7, e33480 (2018).

7. Ward, M. C., Banovich, N. E., Abhishek Sarkar, Stephens, M. & Gilad, Y. Dynamic effects of genetic variation on gene expression revealed following hypoxic stress in cardiomyocytes. eLife 10, e57345 (2021).

8. Cuomo, A. S. E. Single-cell RNA-sequencing of differentiating iPS cells reveals dynamic genetic effects on gene expression. 14 (2020).

9. Pashos, E. E. et al. Large, Diverse Population Cohorts of hiPSCs and Derived Hepatocyte-like Cells Reveal Functional Genetic Variation at Blood Lipid-Associated Loci. Cell Stem Cell 20, 558-570.e10 (2017).

10. Banovich, N. E. et al. Impact of regulatory variation across human iPSCs and differentiated cells. Genome Res. 28, 122–131 (2018).

11. Hung, A., Housman, G., Briscoe, E. A., Cuevas, C. & Gilad, Y. Characterizing gene expression in an in vitro biomechanical strain model of joint health. F1000Research 11, 296 (2022).

12. Kim-Hellmuth, S. et al. Cell type–specific genetic regulation of gene expression across human tissues. Science 369, eaaz8528 (2020).

13. Lancaster, M. A. & Knoblich, J. A. Organogenesis in a dish: Modeling development and disease using organoid technologies. Science 345, 1247125–1247125 (2014).

14. Kim, H., Kamm, R. D., Vunjak-Novakovic, G. & Wu, J. C. Progress in multicellular human cardiac organoids for clinical applications. Cell Stem Cell 29, 503–514 (2022).

15. Thomas, D., Choi, S., Alamana, C., Parker, K. K. & Wu, J. C. Cellular and Engineered Organoids for Cardiovascular Models. Circ. Res. 130, 1780–1802 (2022).

16. Silva, A. C. et al. Co-emergence of cardiac and gut tissues promotes cardiomyocyte maturation within human iPSC-derived organoids. Cell Stem Cell 28, 2137-2152.e6 (2021).

17. Voges, H. K. et al. Vascular cells improve functionality of human cardiac organoids. Cell Rep. 42, 112322 (2023).

18. Bauwens, C. L. et al. Geometric Control of Cardiomyogenic Induction in Human Pluripotent Stem Cells. Tissue Eng. Part A 17, 1901–1909 (2011).

19. Branco, M. A. et al. Transcriptomic analysis of 3D Cardiac Differentiation of Human Induced Pluripotent Stem Cells Reveals Faster Cardiomyocyte Maturation Compared to 2D Culture. Sci. Rep. 9, 9229 (2019).

20. Chen, Y. et al. A versatile polypharmacology platform promotes cytoprotection and viability of human pluripotent and differentiated cells. Nat. Methods 18, 528–541 (2021).

21. Lian, X. et al. Directed cardiomyocyte differentiation from human pluripotent stem cells by modulating Wnt/β-catenin signaling under fully defined conditions. Nat. Protoc. 8, 162–175 (2012).

22. Burridge, P. W. et al. Chemically defined generation of human cardiomyocytes. Nat. Methods 11, 855–860 (2014).

23. Keepers, B., Liu, J. & Qian, L. What’s in a cardiomyocyte – And how do we make one through reprogramming? Biochim. Biophys. Acta BBA - Mol. Cell Res. 1867, 118464 (2020).

24. Lyons, I. et al. Myogenic and morphogenetic defects in the heart tubes of murine embryos lacking the homeo box gene Nkx2-5. Genes Dev. 9, 1654–1666 (1995).

25. Knight-Schrijver, V. R. et al. A single-cell comparison of adult and fetal human epicardium defines the age-associated changes in epicardial activity. Nat. Cardiovasc. Res. 1, 1215–1229 (2022).

26. Acharya, A. et al. The bHLH transcription factor Tcf21 is required for lineage-specific EMT of cardiac fibroblast progenitors. Development 139, 2139–2149 (2012).

27. Jonsson, M. K. B. et al. A Transcriptomic and Epigenomic Comparison of Fetal and Adult Human Cardiac Fibroblasts Reveals Novel Key Transcription Factors in Adult Cardiac Fibroblasts. JACC Basic Transl. Sci. 1, 590–602 (2016).

28. Pinto, A. R. et al. Revisiting Cardiac Cellular Composition. Circ. Res. 118, 400–409 (2016).

29. Ivey, M. J. & Tallquist, M. D. Defining the Cardiac Fibroblast. Circ. J. 80, 2269–2276 (2016).

30. Cui, Y. et al. Single-Cell Transcriptome Analysis Maps the Developmental Track of the Human Heart. Cell Rep. 26, 1934-1950.e5 (2019).

31. Lindqvist, A., Rodríguez-Bravo, V. & Medema, R. H. The decision to enter mitosis: feedback and redundancy in the mitotic entry network. J. Cell Biol. 185, 193–202 (2009).

32. Wang, J. C. DNA topoisomerases. Annu. Rev. Biochem. 54, 665–697 (1985).

33. Tallquist, M. D. & Molkentin, J. D. Redefining the identity of cardiac fibroblasts. Nat. Rev. Cardiol. 14, 484–491 (2017).

34. Kawase, A. et al. Podoplanin expression by cancer associated fibroblasts predicts poor prognosis of lung adenocarcinoma. Int. J. Cancer 123, 1053–1059 (2008).

35. Soldatov, R. et al. Spatiotemporal structure of cell fate decisions in murine neural crest. Science 364, eaas9536 (2019).

36. Sasaki, H. & Hogan, B. L. M. Differential expression of multiple fork head related genes during gastrulation and axial pattern formation in the mouse embryo. Development 118, 47–59 (1993).

37. Mummery, C. et al. Differentiation of Human Embryonic Stem Cells to Cardiomyocytes: Role of Coculture With Visceral Endoderm-Like Cells. Circulation 107, 2733–2740 (2003).

38. Ng, W. H., Varghese, B., Jia, H. & Ren, X. Alliance of Heart and Endoderm: Multilineage Organoids to Model Co-development. Circ. Res. 132, 511–518 (2023).

39. Spence, J. R. et al. Sox17 Regulates Organ Lineage Segregation of Ventral Foregut Progenitor Cells. Dev. Cell 17, 62–74 (2009).

40. Sherwood, R. I., Chen, T. A. & Melton, D. A. Transcriptional dynamics of endodermal organ formation. Dev. Dyn. 238, 29–42 (2009).

41. Öström, M. et al. Retinoic Acid Promotes the Generation of Pancreatic Endocrine Progenitor Cells and Their Further Differentiation into β-Cells. PLoS ONE 3, e2841 (2008).

42. Su, X. et al. Single-cell RNA-Seq analysis reveals dynamic trajectories during mouse liver development. BMC Genomics 18, 946 (2017).

43. Cassia, R. et al. Transferrin is an early marker of hepatic differentiation, and its expression correlates with the postnatal development of oligodendrocytes in mice. J. Neurosci. Res. 50, 421– 432 (1997).

44. Senga, K., Mostov, K. E., Mitaka, T., Miyajima, A. & Tanimizu, N. Grainyhead-like 2 regulates epithelial morphogenesis by establishing functional tight junctions through the organization of a molecular network among claudin3, claudin4, and Rab25. Mol. Biol. Cell 23, 2845–2855 (2012).

45. Kirchhoff, C., Habben, I., Ivell, R. & Krull, N. A Major Human Epididymis-Specific cDNA Encodes a Protein with Sequence Homology to Extracellular Proteinase Inhibitors1. Biol. Reprod. 45, 350– 357 (1991).

46. Du, H., Yang, X., Fan, J. & Du, X. Claudin 6: Therapeutic prospects for tumours, and mechanisms of expression and regulation. Mol. Med. Rep. 24, 677 (2021).

47. Beccari, L., Marco-Ferreres, R. & Bovolenta, P. The logic of gene regulatory networks in early vertebrate forebrain patterning. Mech. Dev. 130, 95–111 (2013).

48. Dominguez, M. H., Ayoub, A. E. & Rakic, P. POU-III Transcription Factors (Brn1, Brn2, and Oct6) Influence Neurogenesis, Molecular Identity, and Migratory Destination of Upper-Layer Cells of the Cerebral Cortex. Cereb. Cortex 23, 2632–2643 (2013).

49. Djuric, U. et al. Spatiotemporal Proteomic Profiling of Human Cerebral Development. Mol. Cell. Proteomics 16, 1548–1562 (2017).

50. Ashique, A. M. et al. The Rfx4 Transcription Factor Modulates Shh Signaling by Regional Control of Ciliogenesis. Sci. Signal. 2, (2009).

51. Southard-Smith, E. M., Kos, L. & Pavan, W. J. SOX10 mutation disrupts neural crest development in Dom Hirschsprung mouse model. Nat. Genet. 18, 60–64 (1998).

52. Goulding, M. D., Chalepakis, G., Deutsch, U., Erselius, J. R. & Gruss, P. Pax-3, a novel murine DNA binding protein expressed during early neurogenesis. EMBO J. 10, 1135–1147 (1991).

53. Knight, R. D. et al. lockjaw encodes a zebrafish tfap2a required for early neural crest development. Development 130, 5755–5768 (2003).

54. Moser, M., Rüschoff, J. & Buettner, R. Comparative analysis of AP-2α and AP-2β gene expression during murine embryogenesis. Dev. Dyn. 208, 115–124 (1997).

55. Schupp, J. C. et al. Integrated Single-Cell Atlas of Endothelial Cells of the Human Lung. Circulation 144, 286–302 (2021).

56. Wu, B. et al. Nfatc1 Coordinates Valve Endocardial Cell Lineage Development Required for Heart Valve Formation. Circ. Res. 109, 183–192 (2011).

57. Zhu, H. et al. Regulation of the lmo2 promoter during hematopoietic and vascular development in zebrafish. Dev. Biol. 281, 256–269 (2005).

58. Lü, J. et al. Isolation and Characterization of EDAG-1, A Novel Gene Related to Regulation in Hematopoietic System. Sheng Wu Hua Xue Yu Sheng Wu Wu Li Xue Bao Acta Biochim. Biophys. Sin. 33, 641–646 (2001).

59. Hoang, T., Lambert, J. A. & Martin, R. SCL/TAL1 in Hematopoiesis and Cellular Reprogramming. in Current Topics in Developmental Biology vol. 118 163–204 (Elsevier, 2016).

60. Palma-Barqueros, V. et al. Platelet transcriptome analysis in patients with germline RUNX1 mutations. J. Thromb. Haemost. 21, 1352–1365 (2023).

61. Macaulay, I. C. et al. Comparative gene expression profiling of in vitro differentiated megakaryocytes and erythroblasts identifies novel activatory and inhibitory platelet membrane proteins. Blood 109, 3260–3269 (2007).

62. Frenis, K. et al. Path of differentiation defines human macrophage identity. Preprint at 10.1101/2025.01.24.634694 (2025).

63. Jeong, R. & Bulyk, M. L. Blood cell traits’ GWAS loci colocalization with variation in PU.1 genomic occupancy prioritizes causal noncoding regulatory variants. Cell Genomics 3, 100327 (2023).

64. Randrianarison-Huetz, V. et al. Gfi-1B controls human erythroid and megakaryocytic differentiation by regulating TGF-β signaling at the bipotent erythro-megakaryocytic progenitor stage. Blood 115, 2784–2795 (2010).

65. Forsberg, E. C. et al. Differential Expression of Novel Potential Regulators in Hematopoietic Stem Cells. PLoS Genet. 1, e28 (2005).

66. Althoff, M. J. et al. Yap1-Scribble polarization is required for hematopoietic stem cell division and fate. Blood 136, 1824–1836 (2020).

67. Dardano, M. et al. Blood-generating heart-forming organoids recapitulate co-development of the human haematopoietic system and the embryonic heart. Nat. Cell Biol. (2024) doi:10.1038/s41556-024-01526-4.

68. Ergen, C. et al. Consensus prediction of cell type labels in single-cell data with popV. Nat. Genet. 56, 2731–2738 (2024).

69. The Tabula Sapiens Consortium et al. The Tabula Sapiens: A multiple-organ, single-cell transcriptomic atlas of humans. Science 376, eabl4896 (2022).

70. The Tabula Sapiens Consortium. Tabula Sapiens reveals transcription factor expression, senescence effects, and sex-specific features in cell types from 28 human organs and tissues. Preprint at 10.1101/2024.12.03.626516 (2024).

71. Umans, B. D., Battle, A. & Gilad, Y. Where Are the Disease-Associated eQTLs? Trends Genet. TIG 37, 109–124 (2021).

72. 20k Human PBMCs, 3’ HT v3.1, Chromium X Single Cell Gene Expression Dataset by Cell Ranger (2021).

73. Huang, Y., McCarthy, D. J. & Stegle, O. Vireo: Bayesian demultiplexing of pooled single-cell RNA-seq data without genotype reference. Genome Biol. 20, 273 (2019).

74. R Core Team. R: A Language and Environment for Statistical Computing. R Foundation for Statistical Computing (2017).

75. RStudio Team. RStudio: Integrated Development Environment for R. RStudio, PBC. (2020).

76. Stuart, T. et al. Comprehensive Integration of Single-Cell Data. Cell 177, 1888-1902.e21 (2019).

77. Wickham, H. et al. Welcome to the Tidyverse. J. Open Source Softw. 4, 1686 (2019).

78. Schindelin, J. et al. Fiji: an open-source platform for biological-image analysis. Nat. Methods 9, 676–682 (2012).

79. Tol, P. Colour Schemes. https://personal.sron.nl/~pault/ (2021).

80. The International HapMap Consortium. The International HapMap Project. Nature 426, 789–796 (2003).

